# Dean flow assisted single cell and bead encapsulation for high performance single cell expression profiling

**DOI:** 10.1101/520858

**Authors:** Luoquan Li, Ping Wu, Zhaofeng Luo, Lei Wang, Weiping Ding, Tao Wu, Jinyu Chen, Jinlong He, Yi He, Heran Wang, Ying Chen, Guibo Li, Zida Li, Liqun He

## Abstract

Single-cell RNA sequencing examines the transcriptome of individual cells and reveals the inter-cell transcription heterogeneity, playing a critical role in both scientific research and clinical applications. Recently, droplet microfluidics-based platform for expression profiling has been shown as a powerful tool to capture of the transcriptional information on single cell level. Despite the breakthrough this platform brought about, it required the simultaneous encapsulation of single cell and single barcoded bead, the incidence of which was very low. Suboptimal capturing efficiency limited the throughput of the Drop-seq platform. In this work, we leveraged the advance in inertial microfluidics-based cell sorting and designed a microfluidic chip for high efficiency cell-bead co-encapsulation, increasing the capturing rate by more than four folds. Specifically, we adopted spiral and serpentine channels and ordered cells/beads before the encapsulation region. We characterized the effect of cell concentration on the capturing rate and achieved a cell-bead co-capturing rate up to 3%. We tested this platform by co-encapsulating barcoded beads and human-mouse cell mixtures. The sequencing data distinguished the majority of human and mice expressions, with the doublet rate being as low as 5.8%, indicating that the simultaneous capturing of two or more cells in one droplet was minimal even when using high cell concentration. This chip design showed great potential in improving the efficiency for future single cell expression profiling.

## 1. Introduction

Conventional biological assays, such as qPCR and western blot, are oftentimes performed on bulk tissue level. Such assaying strategies assumed the overall tissue as a homogeneous entity and gauged the corresponding average values. Recently, it is more and more appreciated that the heterogeneity of individual cells within a tissue has ineligible impact on the analysis of assay results^1-2^. Indeed, single cell analysis has played a significant role in discovering unseen biological mechanisms as well as in making informed and personalized treatment plans in clinics. Heterogeneous gene structure and gene expression status of individual cells have been revealed on single cell level by whole genome sequencing^3^, transcriptome sequencing^4-5^, and epigenetic sequencing^6^.

Single-cell transcriptome sequencing (scRNA-seq) gained great attention in recent years as a method for studying individual cell transcriptomes on large scale^7-9^. It revealed in detail the gene expression of whole tissues and elucidated the genetic and epigenetic mechanisms. Compared to conventional assays performed on bulk tissue level, scRNA-seq can quantify intrapopulation heterogeneity and dissect the transcriptional landscape of single cells at very high resolution. In addition, scRNA-seq is important in the evaluation of genetically heterogeneous tumors, significantly impacting the battle against cancer.

Microfluidics is a technology that provides exquisite flow control with low sample consumption. Thanks to the advantages that microfluidics possesses, researchers developed high-throughput single-cell analysis platforms based on this technology^10-13^. Microfluidic chip can achieve high throughput, high separation efficiency, and low sample demand, which is ideal for automated sample preparation and single cell assays. Various devices can also be integrated on one chip, making the chip highly versatile. In 2015, Macosko et al. reported a high-throughput scRNA-seq method based on droplet microfluidics (Drop-seq)^10^, which encapsulated a single cell along with a single barcoded bead into nanoliter droplets. The barcode of each bead, appearing in the sequencing data, served as the identifier of individual cells and helped trace the origin of each read. This smart design significantly reduced the bias in PCR amplification and noise in sequencing data analysis.

Despite the great potential that Drop-seq has shown, this platform has major challenges. The nature of the experiment required that a single cell and a single bead being simultaneously encapsulated in a single droplet and strictly prohibited encapsulation of multiple cells. The encapsulation incidence followed Poisson distribution; to minimize multiple encapsulation, the cell/bead suspension needed to highly diluted. As a result, the encapsulation rate was reported to be as low as 0.15% under low cell/bead concentration of 100 μL^-110^. The extremely low encapsulation rate inevitably compromised the efficiency and the throughput of the platform.

Inertial focusing provides a way to hydrodynamically localize particles in certain regions of the channel cross section. Consequently, particles line up and approach the encapsulation region one by one, thus reducing the incidence of multiple encapsulation and improving the encapsulation efficiency. When particles flow in microfluidic channels, due to the parabolic velocity profile, they are subject to lift forces, namely shear gradient lift force and wall effect lift force. In curved channels, the inertia of the fluids generates secondary swirling flow, named Dean flow, which accelerates the lateral displacement of the particles and thus facilitates the focusing. Based on the principle of inertial focusing, spiral channels were adopted in a reported work to order barcoded beads before droplet encapsulation^14^. Higher capturing efficiency and higher fraction of single bead encapsulation were reported.

Despite the efficient focusing that has been achieved for beads, cell focusing has been challenging, partly due to that lift force scales with the fourth power of particle diameter^15^. Cells, usually more than twice smaller than beads, take much longer to be focused in spiral channels. With efficient cell ordering, the cell/bead co-encapsulating rate could be further enhanced. To this end, here we developed a new microfluidic chip which inertially ordered both beads and cells, aiming for high throughput single cell expression profiling. To efficiently focus cells, we coupled serpentine channels with spiral channels before the droplet generation region. Consequently, the incidence of multiple cell encapsulation was greatly reduced even at high cell/bead concentrations, which maximized the sample utilization and increased the throughput. We investigated the effect of cell/bead concentration and size on the encapsulation performance. We further tested the utility of this platform by performing single cell sequencing of human-mouse cell mixture and successfully distinguished nearly 95% of the two populations in the sequencing data, suggesting that the incidence of multiple encapsulation was sufficiently low. Our platform showed the great potential for single cell expression profiling with improved throughput.

## 2. Materials and methods

### 2.1. Bead preparation

Plain beads without surface functionalization (CM-300-10 and CM-100-10, Spherotech Inc., USA) with diameters of 10 μm and 30 μm were used in lieu of cells and barcoded beads, respectively, to test encapsulation efficiencies. The density of the bead phase solution was adjusted by supplementing OptiPrep Density Gradient Medium (D1556, Sigma Aldrich, USA) to alleviate bead sedimentation.

Barcoded beads (OSKO-2011-10, ChemGenes) functionalized with millions of primers were used to capture mRNA released from lysed cells in the Drop-seq experiment. The primers on one bead consist of a common PCR sequence, an identical cell barcode, variety of unique molecular identifiers (UMIs), and poly-T tail. Upon delivery, the beads were first gently washed twice with 100% ethanol to remove free nucleotide sequences, before being washed and resuspended in TE/TW buffer (TE Buffer pH 8.0, 0.01% Tween). The suspension was then filtered through a 100 μm filter (BD Falcon) to remove aggregates and the filtrates were aliquoted into multiple tubes for future use. Beads were stored strictly between 3.5 and 4 °C. Before each experiment, the beads were resuspended in lysis buffer containing 200 mM Tris, pH 7.5, 6% Ficoll PM-400 (17-0300-10, GE Healthcare, Sweden), 0.2% sarcosyl (L7414, Sigma, USA), and 20 mM EDTA. Dithiothreitol was supplemented to the lysate prior to droplet generation.

### 2.2. Cell preparation

Two cell lines (Human HEK293T and Mouse NIH3T3) were used. Cells were cultured in high-glucose Dulbecco’s Modified Eagle’s medium (DMEM; Gibco, Fisher Scientific) supplemented with 10% (v/v) fetal bovine serum (Gibco, Fisher Scientific) and 1% penicillin-streptomycin antibiotics (Invitrogen, USA). Cells were passaged at a confluency of about 75%.

### 2.3. Microfluidic devices

The chip design adopted spiral channels following the inlets to focus cells/beads, as shown in Figure 1a. The microfluidic chip was fabricated with standard soft lithography and the mold was fabricated using SU-8 photolithography. Polydimethylsiloxane (PDMS; Sylgard 184, Dow Corning, USA) with 10:1 base-to-curing agent ratio was poured on the mold and degassed for about 10 minutes, before it was baked in 80 °C oven for 40 minutes to cure. After baking, the PDMS was peeled off from the wafer and cut with razor blades. Biopsy punches were used to punch holes for tubing connection. Finally, the PDMS slab was bonded to glass slide after surface activation with oxygen plasma (Harrick Plasma, USA) followed by another baking at 80 °C. The channels were rendered hydrophobic by treating with Aquapel.

**Figure 1.**
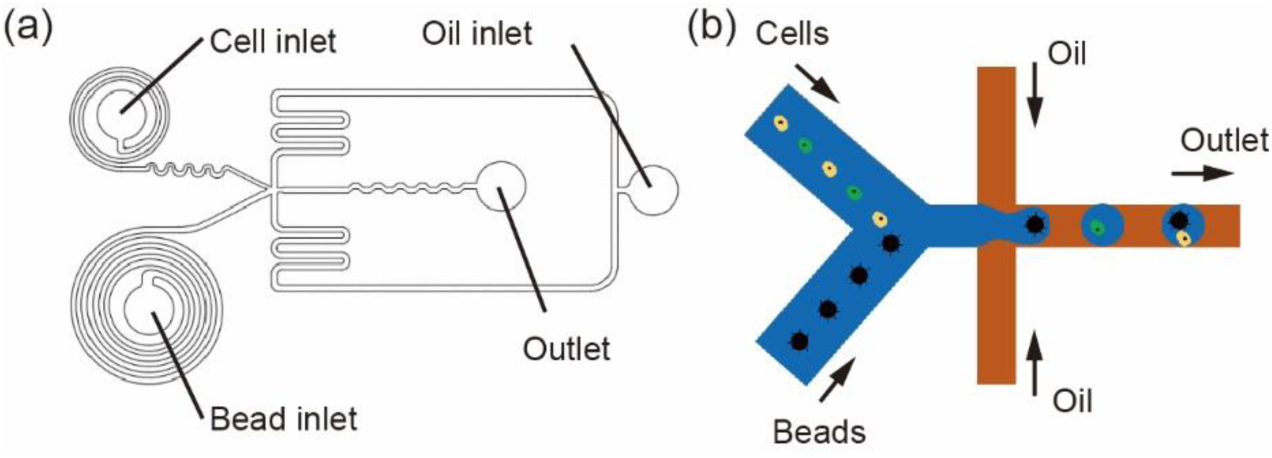
Experimental setup. (a) Schematic of the design of the microfluidic chip. Spiral channels were adopted for cell and bead ordering. Due to the small size of cells, serpentine channels were implemented to enhance the performance of cell ordering. Cell suspension, bead suspensions, and carrier oil were infused into the chip from inlets as indicated. Droplets were collected from the outlet. (b) Schematic showing droplet generation and bead/cell encapsulation.

The chip design incorporated three inlets, namely cell, bead, and carrier oil inlet, and one outlet. The chip had three functional units: two spiral structures with serpentine channels for cell/bead ordering, a droplet generator, and a serpentine channel for efficient mixing of generated droplets. The chip had a height of 120 μm throughout. Cell channel was a 2-loop spiral channels with a width of 80 μm. The innermost layer has a radius of 740 μm, and the interval between adjacent two loops was 100 μm. The spiral channel was followed by an asymmetric serpentine channel. Bead channel was 5-loop spiral channels with a radius of 760 μm for the innermost layer, 100 μm intervals between adjacent loops, and 120 μm channel width.

### 2.4. Droplet generation and cell/bead encapsulation

Negative pressure was used to drive the flows in the microfluidic chip. Cells were suspended in 1x PBS supplemented with 0.01% BSA (v/v) and beads were suspended in lysis agent. The bead/cell suspensions were stored in 1.5 mL centrifuge tubes and connected to the chip via polyethylene tubing (BB31695-PE/2, Scientific Commodities Inc.). Droplets were collected in a customized centrifuge tube with inlet and outlet on the lid. The outlet was connected to a syringe pump, which was set on withdrawal mode with a typical flow rate of 30 mL/h to generate the negative pressure and form droplets stably^16-17^. An inverted microscope (IX71, Olympus, Germany), coupled with a high-speed camera (DP26, Olympus, Germany) was used to acquire images. ImageJ (National Institute of Health, USA) was used to analyze the acquired images.

As soon as the cell was captured in the droplet, it was rapidly lysed by the lysis buffer in the bead phase and primers on the barcoded bead captured messenger RNAs (mRNA). The collected droplets were broken by adding perfluoro octanol (370533-5G, Sigma), and the beads were washed with large volume of 6x saline sodium citrate buffer (SSC) and harvested. The mRNAs were reverse transcribed together to form stable single-cell transcriptomes bonded to beads and then amplified.

### 2.5. Library construction and sequencing

Overall, qualified library was built after completion of DNA fragmentation and cyclization. We used dsDNA fragmentase (M0348L, NEB) to interrupt the target sequence (cDNA) randomly, followed by end-repairing and adding adapter primers^18^. The fragmented cDNA was amplified with two specific primers. Subsequently, DNA mix was selected to 300-500 bp by AMPure XP beads (Beckman). Finally, the DNA was heat denatured and cyclized into single-strand circular DNA with the aid of exogenous free single-stranded nucleotide primers. We made DNA nanoball using rolling circle amplification^19-20^ and sequenced the resultant molecules from each end using BGISEQ-500 sequencer (MGI Tech Co., China). The detailed library construction procedures can be found in Fig. S1 and the relevant oligonucleotide sequences in the Drop-seq experiments can also be found in Table S3.

### 2.6. Analysis of the scRNA-seq data

**3.** With the obtained paired-end sequencing data, we first convert the data to sam/bam format using Picard (v2.9.3). We applied Drop-seq tools v1.13^10^ to decode UMI and cell barcode information embedded in the sequences in order to quantify the transcript abundance. The UMI and cell barcode encoded in Read 1 (forward strand) sequences are abstracted after filtering out low-quality sequences. As for Read 2 (reverse strand) sequences, which are 100 bp mRNA insert sequences, we removed the sequences with high adapter and poly A contamination and used STAR (v2.5.2b) to map the read sequences to a reference for gene alignment^9^. After quantifying the UMI counts for each cell barcode, we selected cells that have higher number of UMIs (evaluated by ‘expect cell numbers’ or a single ‘UMI cut-off’) as the valid cells.**Results and discussion**

### 3.1. Overall experimental design

This chip aimed to focus cells/beads before droplet generation and encapsulation and improve the encapsulation efficiency. When cells/beads are focused, they would align as a train of particles and enter the droplet generating region one by one, reducing the chance of multiple encapsulation.

The microfluidic device adopted spiral and serpentine channels to focus the cells/beads. In spiral channels, due to the viscosity of fluid and velocity imbalance between the channel center and the near-wall region, secondary flow, namely Dean flow, was generated, leading to Dean drag force on the particles. In addition, particles were also subject to lift force, which was the net force of the inertial lift, caused by the shear gradient, and the wall lift, caused by the channel wall^21-22^. The balance of drag force and lift force dictated the equilibrium positions of the particles on the channel cross section^14^. Inertial microfluidics predicted that particles are more likely to occupy a single equilibrium position in a curvilinear channel when^21-25^

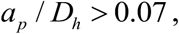

Where *a*_*p*_ is the diameter of particles and *D*_*h*_ is the hydraulic diameter. Additionally, the net lift force, *F*_*L*_, can be estimated by^21-22, 26^

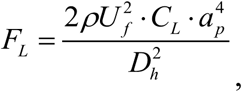

Where *ρ* is the density of the fluid, *U* _*f*_ is the flow velocity of the fluid, *C*_*L*_ is the lift coefficient, which is a function of the particle position across the cross-section of the microchannel and has an average value of 0.5. Given that particle mass scales with *a*_*p*_ ^3^ and *F*_*L*_ scales with *a*_*p*_ ^4^, smaller particles, such as cells, would experience lower acceleration and take longer time to reach the equilibrium position. Combined together, designs of spiral channels would be sufficient to focus beads, but for cells, we added serpentine channels^15^ at the outlet of the spiral channel as an additional focusing mechanism, as shown in Figure 1a. In the asymmetrical semicircle serpentine channel, cells were simultaneously subjected to both viscous drag force (*F*_*D*_) and centrifugal force (*F*_*C*_), and the equilibrium position of cells depends on the balance of the two forces^15, 27^. As cells and beads approached the droplet generation region (Figure 1b), they were encapsulated into droplets. Since the cells/beads were focused on a lateral position, the proportion of multiple encapsulation in single droplets were reduced. Cell lysis buffer loaded in the bead suspension lysed the cell, exposing its mRNA to the primers functionalized on the bead surface which captured the poly-A tail of the mRNA (see Methods).

We examined the performance of bead/cell ordering at different bead/cell concentrations. Due to their larger sizes, beads were well focused at lateral positions close to the outer channel walls at concentrations ranging from 250 to 1000 μL^-1^, as shown in Figure 2. However, we noticed that at high concentrations, namely 1000 μL^-1^, the spacing between beads became small, increasing the chance of getting multiple encapsulations. We also achieved cell focusing with the assistance of serpentine channels at cell concentrations of 450 and 900 μL^-1^.

**Figure 2.**
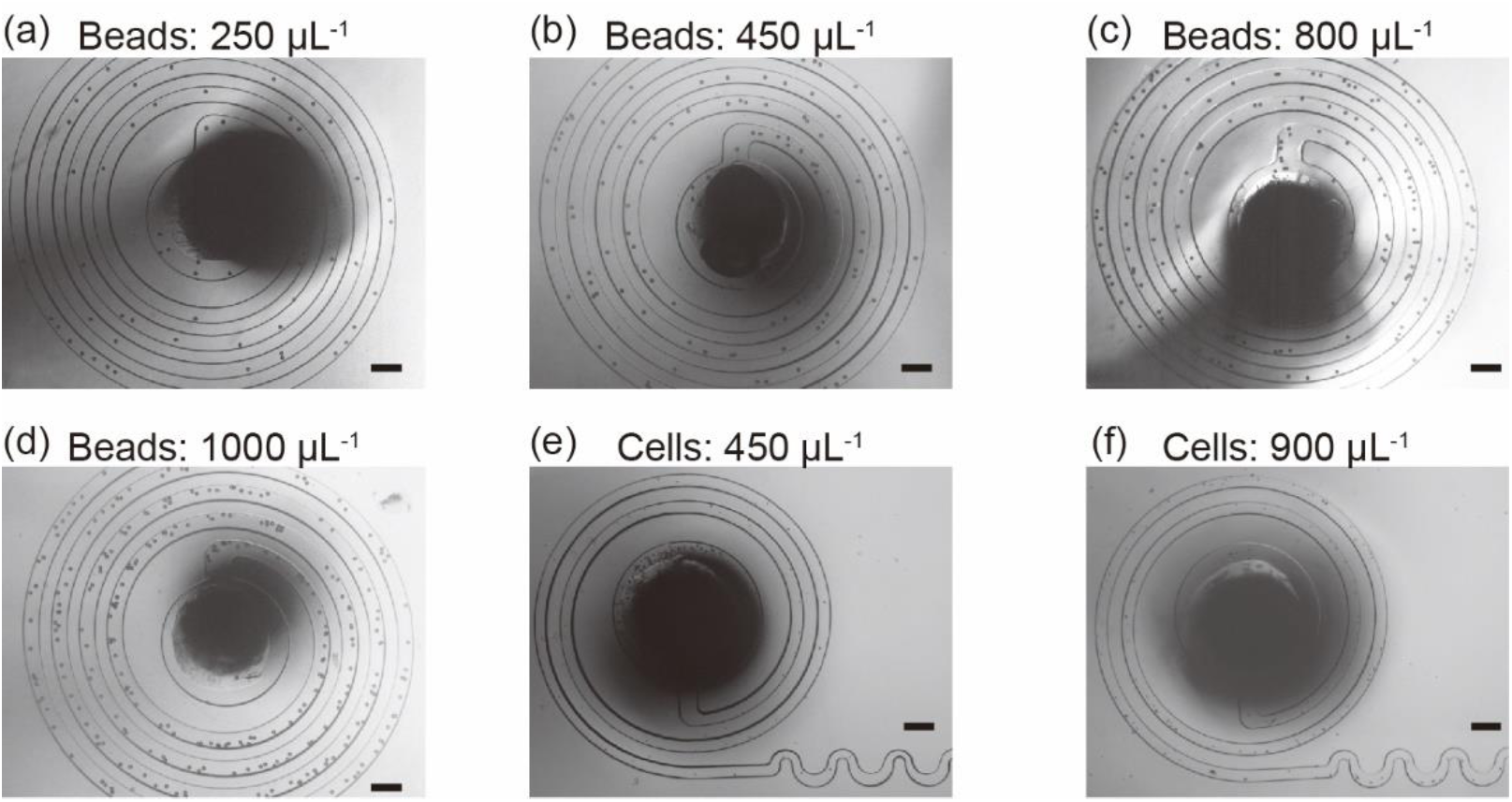
Photographs showing the bead and cell ordering at different concentrations. Beads flowed in five-loop spiral channels and cells flowed in the two-loop spiral channels followed by serpentine channels. (a-d) Bead ordering at different bead concentrations as indicated. Beads ordered with even spacing when the bead concentration varied from 250 to 800 μL^-1^. Higher bead concentration led to uneven spacing while the ordering was not significantly affected. (e & f) Cell ordering at different cell concentrations as indicated. Scale bars: 200 μm.

### 3.2. Single bead and single cell encapsulation rates

To achieve cell/bead co-encapsulation with high efficiency, it was necessary to achieve cell/bead encapsulation with high efficiency. Thus, we investigated the encapsulation of beads/cells separated.

Plain beads with a diameter of 30 μm were used to test the bead encapsulation efficiency. Intuitively, the fraction of single encapsulation in the resultant droplets would increase as the bead suspension became more concentrated, with a tradeoff that multiple encapsulation would increase as well. Indeed, as shown in Figure 3a & b, we observed that when the bead concentration was lower than 900 μL^-1^, we were able to achieve a single encapsulation rate up to 23.97%. As the bead concentration went higher, we started to see encapsulations with multiple beads in single droplets. For example, at concentration of 1100 μL^-1^, while the fraction of single bead encapsulation was increased to 29.83%, we also observed 2.86% droplets with multiple beads encapsulated. Encapsulation with multiple beads lead to false sequencing data; therefore, bead concentration lower than 900 μL^-1^ was used in the following experiments. In the test experiments of cell encapsulation, we obtained similar observations (Figure 3c & d). Cell concentration at 700 μL^-1^ resulted in a single cell encapsulation rate of 19.7% with 1.14% multiple cell encapsulation. Therefore, cell concentrations lower than 700 μL^-1^ was adopted. The detailed experimental data can be found in Table S1 & S2 in Supplementary Information.

**Figure 3.**
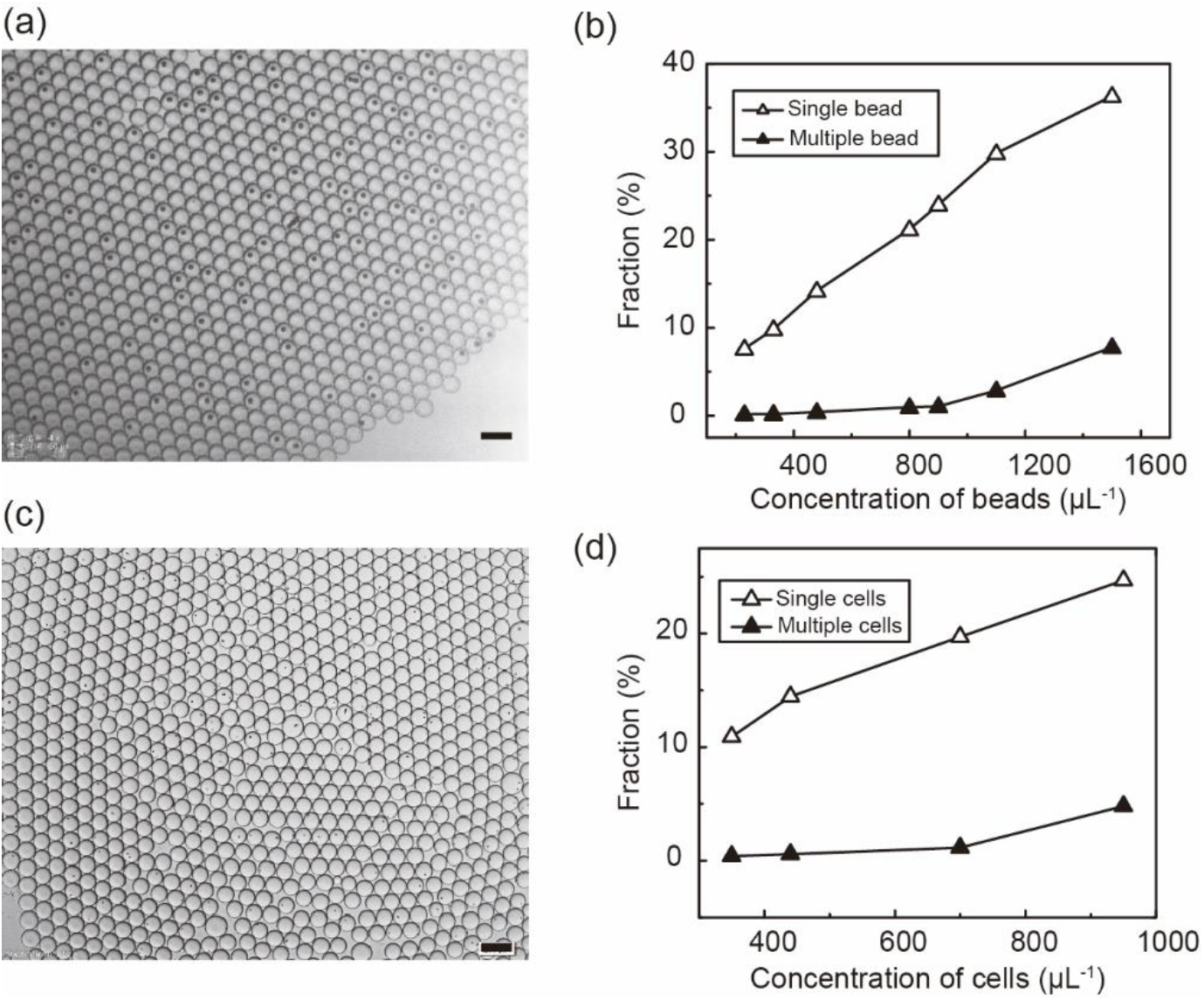
Performance of bead/cell encapsulation at different bead/cell concentrations. (a) A micrograph showing droplets with encapsulated beads. (b) Fraction of droplets encapsulated with single or multiple beads at different bead concentrations. (c) A Photograph showing droplets with encapsulate cells. (d) Fraction of droplets encapsulated with single or multiple cells at different cell concentrations. Scale bars, 200 μm.

### 3.3. Bead and cell co-encapsulation

We loaded both bead and cell suspensions into the device and tested the cell/bead co-encapsulation performance. Compared to the microfluidic design with no spiral designs, where cells were present at multiple the lateral positions, in this proposed design, beads and cells lined up and entered the droplet generation region one by one, which facilitated single encapsulation (Figure 4a & b). At bead concentration of 900 μL^-1^ and cell concentration of 300 μL^-1^, the co-encapsulation rate of a bead and a cell was about 3% with very low cell doublet or bead doublet rate (Figure 4c & d). Thousands of effective encapsulations were generated within minutes.

**Figure 4.**
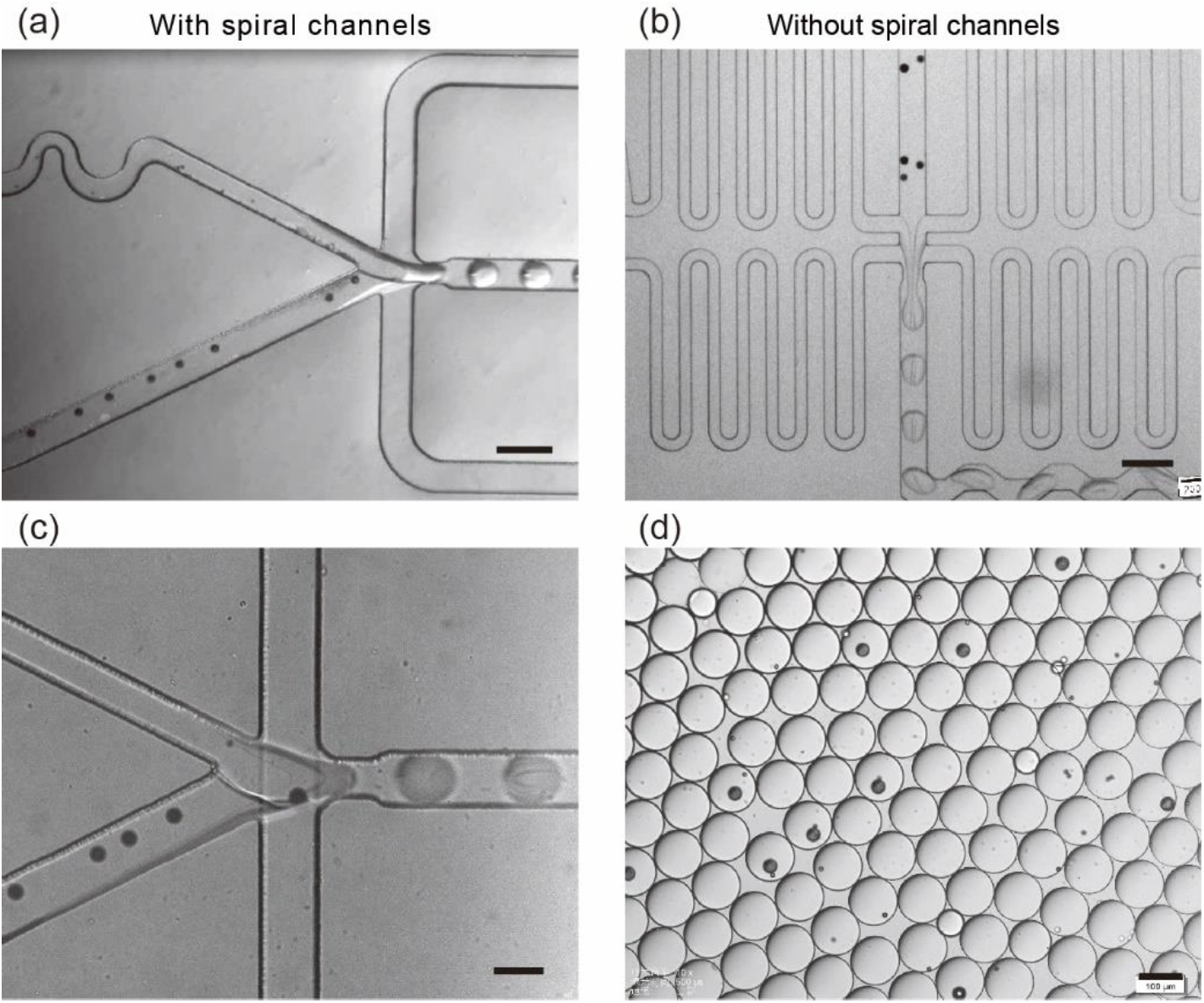
(a & b) Comparison of bead and cell ordering before encapsulation region with and without spiral and serpentine channels, as indicated. Scale bars, 200 μm. (c) A zoom-in micrograph showing the cell/bead encapsulation process. Scale bar, 200 μm. (d) A micrograph showing the resultant droplets. Scale bar, 100 μm.

### 3.4. scRNA-seq with human and mouse cell mixture

To validate the utility of this microfluidic platform, we used cell mixtures from two cell lines, namely human HEK293T and mouse NIH3T3, and performed cell/bead co-encapsulation and downstream expression profiling. Based on the results we obtained in the aforementioned testing experiments, cell suspensions were loaded at concentration of 300 μL^-1^ and barcoded beads at 900 μL^-1^. Upon encapsulation, cells were lysed and mRNA were captured by the barcoded beads. Droplets were collected into a tube for 10 minutes and then coalesced, and the mRNAs captured on the barcoded beads were subsequently reverse transcribed and amplified. Finally, library was constructed and sequenced on BGISEQ-500.

Each sequence (transcript) was mapped to human/mouse reference genome, as shown in Figure 5a. The species of the analyzed cells was classified if 90% of transcripts were mapped to either of the species. Cells that had both human and mouse transcripts were regarded as cell doublet, since it indicated that the primers on the bead simultaneously captured mRNAs from both a human cell and a mouse cell.

**Figure 5.**
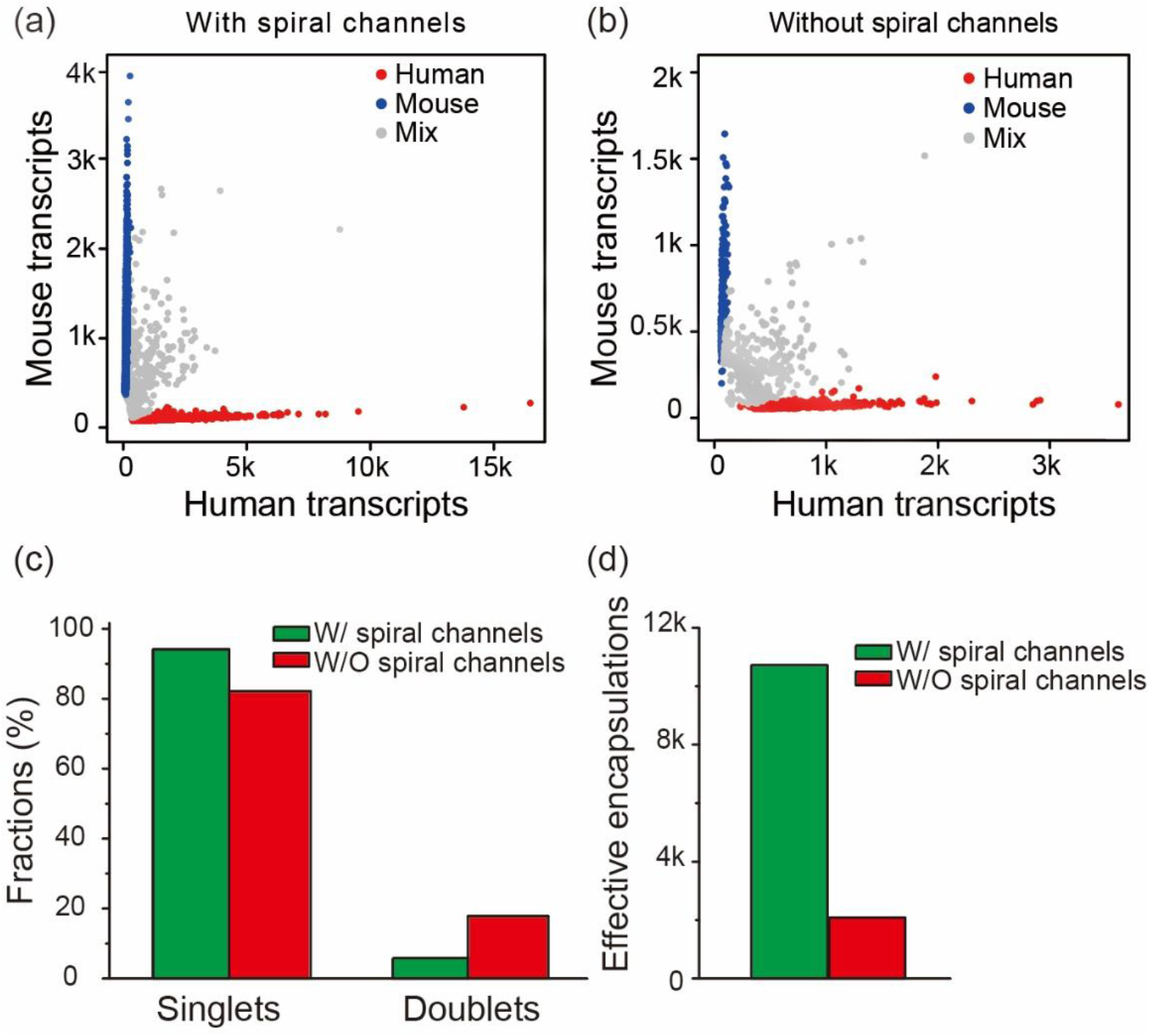
Comparison of the sequencing data obtained from devices with spiral channels and without spiral channels. (a & b) Species plot based on the analyzed transcripts from the sequencing results. Each scattered dot represents the read from a barcoded bead, being classified as single human cell, single mouse cell, or doublet, as indicated. The loaded concentration of cells and beads were 300 and 900 μL^-1^, respectively. (c) Bar plot of the fractions of singlets and doublets analyzed from the sequencing data. (d) Bar plot of the number of effective encapsulations.

We compared sequencing results from this microfluidic design with spiral channels and the reported microfluidic design in Drop-seq without spiral channels. Both experiments adopted the same flow rates and cell and bead concentrations. Based on the sequencing data, the majority of the effective encapsulations captured either a single human cell or a single mouse cell, as shown in Figure 5a & b. Using the design with spiral channels, 94.19% of the effective encapsulations captured either a single human cell or a single mouse cell, with only 5.81% doublets. Similarly, with the design without spiral channels, the fraction of single cell encapsulation was as high as 82.18%. However, the fraction of doublets was 17.82%, nearly three folds higher than that of design with spiral channels (Figure 5c). Notice that the design with spiral channels generated more effective encapsulations, namely 10719, which was about five folds more than that without spiral channels (Figure 5d). These results indicated that the design with spiral and serpentine channels could greatly improve the encapsulation efficiency.

## 4. Conclusion

In this work, we presented a microfluidic design with improved performance for single-cell expression profiling. We adopted spiral and serpentine channels to focus beads and cells, resulting in ordered beads and cells flows, which reduced the incidence of multiple encapsulation. Results showed that the fraction of multiple beads encapsulation was less than 1% and the fraction of multiple cell encapsulation was less than 5%, which were much smaller than the results from Drop-seq without inertial ordering, namely 10% and 20%, respectively. In order to verify the practical utility of this microfluidic platform, we conducted experiments using mixtures of human and mouse cells and performed single-cell sequencing. The results confirmed the capability of this chip design in throughput improvement, which would ensure maximal utilization of the cells to be analyzed. This platform would be extremely useful in single cell analysis where samples are scarce.

## Data Availability

The sequencing raw data reported in this study have been deposited to CNGB Nucleotide Sequence Archive (Accession Code: CNP0000223).

## Acknowledgements

This study was supported by the National Natural Science Foundation of China (Grant No. 31500694, 31670866, and 31570755) and Shenzhen Peacock Talent Plan (No. KQTD20150330171505310).

## References

1. Meacham, C. E.; Morrison, S. J., Tumour heterogeneity and cancer cell plasticity. Nature 2013, 501 (7467), 328–37.

2. Aw Yong, K. M.; Li, Z.; Merajver, S. D.; Fu, J., Tracking the tumor invasion front using long-term fluidic tumoroid culture. Sci Rep 2017, 7 (1), 10784.

3. Mardis, E. R., Next-generation DNA sequencing methods. Annu Rev Genomics Hum Genet 2008, 9, 387–402.

4. Wu, A. R.; Neff, N. F.; Kalisky, T.; Dalerba, P.; Treutlein, B.; Rothenberg, M. E.; Mburu, F. M.; Mantalas, G. L.; Sim, S.; Clarke, M. F.; Quake, S. R., Quantitative assessment of single-cell RNA-sequencing methods. Nat Methods 2014, 11 (1), 41–6.

5. Hashimshony, T.; Senderovich, N.; Avital, G.; Klochendler, A.; de Leeuw, Y.; Anavy, L.; Gennert, D.; Li, S.; Livak, K. J.; Rozenblatt-Rosen, O.; Dor, Y.; Regev, A.; Yanai, I., CEL-Seq2: sensitive highly-multiplexed single-cell RNA-Seq. Genome Biol 2016, 17, 77.

6. Harris, R. A.; Wang, T.; Coarfa, C.; Nagarajan, R. P.; Hong, C.; Downey, S. L.; Johnson, B. E.; Fouse, S. D.; Delaney, A.; Zhao, Y.; Olshen, A.; Ballinger, T.; Zhou, X.; Forsberg, K. J.; Gu, J.; Echipare, L.; O’Geen, H.; Lister, R.; Pelizzola, M.; Xi, Y.; Epstein, C. B.; Bernstein, B. E.; Hawkins, R. D.; Ren, B.; Chung, W. Y.; Gu, H.; Bock, C.; Gnirke, A.; Zhang, M. Q.; Haussler, D.; Ecker, J. R.; Li, W.; Farnham, P. J.; Waterland, R. A.; Meissner, A.; Marra, M. A.; Hirst, M.; Milosavljevic, A.; Costello, J. F., Comparison of sequencing-based methods to profile DNA methylation and identification of monoallelic epigenetic modifications. Nat Biotechnol 2010, 28 (10), 1097–105.

7. Dal Molin, A.; Di Camillo, B., How to design a single-cell RNA-sequencing experiment: pitfalls, challenges and perspectives. Brief Bioinform 2018.

8. Xu, Y.; Zhou, X., Applications of Single-Cell Sequencing for Multiomics. Methods Mol Biol 2018, 1754, 327–374.

9. Hwang, B.; Lee, J. H.; Bang, D., Single-cell RNA sequencing technologies and bioinformatics pipelines. Exp Mol Med 2018, 50 (8), 96.

10. Macosko, E. Z.; Basu, A.; Satija, R.; Nemesh, J.; Shekhar, K.; Goldman, M.; Tirosh, I.; Bialas, A. R.; Kamitaki, N.; Martersteck, E. M.; Trombetta, J. J.; Weitz, D. A.; Sanes, J. R.; Shalek, A. K.; Regev, A.; McCarroll, S. A., Highly Parallel Genome-wide Expression Profiling of Individual Cells Using Nanoliter Droplets. Cell 2015, 161 (5), 1202–1214.

11. Klein, A. M.; Mazutis, L.; Akartuna, I.; Tallapragada, N.; Veres, A.; Li, V.; Peshkin, L.; Weitz, D. A.; Kirschner, M. W., Droplet barcoding for single-cell transcriptomics applied to embryonic stem cells. Cell 2015, 161 (5), 1187–1201.

12. Hosokawa, M.; Nishikawa, Y.; Kogawa, M.; Takeyama, H., Massively parallel whole genome amplification for single-cell sequencing using droplet microfluidics. Sci Rep 2017, 7 (1), 5199.

13. Habib, N.; Avraham-Davidi, I.; Basu, A.; Burks, T.; Shekhar, K.; Hofree, M.; Choudhury, S. R.; Aguet, F.; Gelfand, E.; Ardlie, K.; Weitz, D. A.; Rozenblatt-Rosen, O.; Zhang, F.; Regev, A., Massively parallel single-nucleus RNA-seq with DroNc-seq. Nat Methods 2017, 14 (10), 955–958.

14. Moon, H. S.; Je, K.; Min, J. W.; Park, D.; Han, K. Y.; Shin, S. H.; Park, W. Y.; Yoo, C. E.; Kim, S. H., Inertial-ordering-assisted droplet microfluidics for high-throughput single-cell RNA-sequencing. Lab Chip 2018, 18 (5), 775–784.

15. Wang, L.; Dandy, D. S., High-Throughput Inertial Focusing of Micrometer- and Sub-Micrometer-Sized Particles Separation. Adv Sci (Weinh) 2017, 4 (10), 1700153.

16. Li, Z.; Mak, S. Y.; Sauret, A.; Shum, H. C., Syringe-pump-induced fluctuation in all-aqueous microfluidic system implications for flow rate accuracy. Lab Chip 2014, 14 (4), 744–9.

17. Mak, S. Y.; Li, Z.; Frere, A.; Chan, T. C.; Shum, H. C., Musical interfaces: visualization and reconstruction of music with a microfluidic two-phase flow. Sci Rep 2014, 4, 6675.

18. Huang, J.; Liang, X.; Xuan, Y.; Geng, C.; Li, Y.; Lu, H.; Qu, S.; Mei, X.; Chen, H.; Yu, T.; Sun, N.; Rao, J.; Wang, J.; Zhang, W.; Chen, Y.; Liao, S.; Jiang, H.; Liu, X.; Yang, Z.; Mu, F.; Gao, S., A reference human genome dataset of the BGISEQ-500 sequencer. Gigascience 2017, 6 (5), 1–9.

19. Drmanac, R.; Sparks, A. B.; Callow, M. J.; Halpern, A. L.; Burns, N. L.; Kermani, B. G.; Carnevali, P.; Nazarenko, I.; Nilsen, G. B.; Yeung, G.; Dahl, F.; Fernandez, A.; Staker, B.; Pant, K. P.; Baccash, J.; Borcherding, A. P.; Brownley, A.; Cedeno, R.; Chen, L.; Chernikoff, D.; Cheung, A.; Chirita, R.; Curson, B.; Ebert, J. C.; Hacker, C. R.; Hartlage, R.; Hauser, B.; Huang, S.; Jiang, Y.; Karpinchyk, V.; Koenig, M.; Kong, C.; Landers, T.; Le, C.; Liu, J.; McBride, C. E.; Morenzoni, M.; Morey, R. E.; Mutch, K.; Perazich, H.; Perry, K.; Peters, B. A.; Peterson, J.; Pethiyagoda, C. L.; Pothuraju, K.; Richter, C.; Rosenbaum, A. M.; Roy, S.; Shafto, J.; Sharanhovich, U.; Shannon, K. W.; Sheppy, C. G.; Sun, M.; Thakuria, J. V.; Tran, A.; Vu, D.; Zaranek, A. W.; Wu, X.; Drmanac, S.; Oliphant, A. R.; Banyai, W. C.; Martin, B.; Ballinger, D. G.; Church, G. M.; Reid, C. A., Human genome sequencing using unchained base reads on self-assembling DNA nanoarrays. Science 2010, 327 (5961), 78–81.

20. Fehlmann, T.; Reinheimer, S.; Geng, C.; Su, X.; Drmanac, S.; Alexeev, A.; Zhang, C.; Backes, C.; Ludwig, N.; Hart, M.; An, D.; Zhu, Z.; Xu, C.; Chen, A.; Ni, M.; Liu, J.; Li, Y.; Poulter, M.; Li, Y.; Stahler, C.; Drmanac, R.; Xu, X.; Meese, E.; Keller, A., cPAS-based sequencing on the BGISEQ-500 to explore small non-coding RNAs. Clin Epigenetics 2016, 8, 123.

21. Ramachandraiah, H.; Ardabili, S.; Faridi, A. M.; Gantelius, J.; Kowalewski, J. M.; Martensson, G.; Russom, A., Dean flow-coupled inertial focusing in curved channels. Biomicrofluidics 2014, 8 (3), 034117.

22. Kemna, E. W.; Schoeman, R. M.; Wolbers, F.; Vermes, I.; Weitz, D. A.; van den Berg, A., High-yield cell ordering and deterministic cell-in-droplet encapsulation using Dean flow in a curved microchannel. Lab Chip 2012, 12 (16), 2881–7.

23. Hasni, A. E.; Göb bels, K.; Thiebes, A. L.; Bräunig, P.; Mokwa, W.; Schnakenberg, U., Focusing and Sorting of Particles in Spiral Microfluidic Channels. Procedia Engineering 2011, 25, 1197–1200.

24. Nivedita, N.; Ligrani, P.; Papautsky, I., Dean Flow Dynamics in Low-Aspect Ratio Spiral Microchannels. Sci Rep 2017, 7, 44072.

25. Xiang, N.; Ni, Z.; Yi, H., Concentration-controlled particle focusing in spiral elasto-inertial microfluidic devices. Electrophoresis 2018, 39 (2), 417–424.

26. Bhagat, A. A. S.; Kuntaegowdanahalli, S. S.; Papautsky, I., Inertial microfluidics for continuous particle filtration and extraction. Microfluidics and Nanofluidics 2008, 7 (2), 217–226.

27. Di Carlo, D.; Irimia, D.; Tompkins, R. G.; Toner, M., Continuous inertial focusing, ordering, and separation of particles in microchannels. Proc Natl Acad Sci U S A 2007, 104 (48), 18892–7.

