## Supplementary Information for "Dean flow assisted single cell and bead encapsulation for high performance single cell expression profiling"

^a^Department of Thermal Science and Energy Engineering, University of Science and Technology of China, Hefei 230027, China; ^b^BGI-Shenzhen, Shenzhen 518083, China; ^c^China National GeneBank, BGI-Shenzhen, Shenzhen 518120, China; ^d^School of Life Sciences, University of Science and Technology of China, Hefei 230027, China; ^e^Department of Electronic Science and Technology, University of Science and Technology of China, Hefei 230027, China; ^f^School of Materials and Energy, Guangdong University of Technology, Guangzhou 510006, China; ^g^Department of Biomedical Engineering, School of Medicine, Shenzhen University, Shenzhen 518060, China; ^h^Hefei Energy Research Institute, Hefei 230051, China.

#Equal contribution.

**Table S1**. Encapsulation rates at different bead concentrations.

| Concentration of beads (μL^-1^) | Total amount of droplets | Amount of droplets containing a single bead | Fraction of droplets containing a single bead (%) | Amount of droplets containing multiple beads | Fraction of droplets containing multiple beads (%) |
| --- | --- | --- | --- | --- | --- |
| 230 | 16386 | 1250 | 7.63 | 28 | 0.17 |
| 330 | 4890 | 481 | 9.84 | 10 | 0.20 |
| 480 | 3566 | 566 | 14.19 | 15 | 0.42 |
| 800 | 6120 | 1297 | 21.19 | 59 | 0.96 |
| 900 | 5535 | 1327 | 23.97 | 60 | 1.08 |
| 1100 | 15055 | 4491 | 29.83 | 430 | 2.86 |
| 1500 | 5325 | 1935 | 36.34 | 416 | 7.80 |

**Table S2.** Encapsulation rates at different cell concentrations.

| Concentration of cells (μL^-1^) | Total number of droplets | Amount of droplets containing a single cell | Fraction of droplets containing a single cell (%) | Amount of droplets containing multiple cells | Fraction of droplets containing multiple cells (%) |
| --- | --- | --- | --- | --- | --- |
| 340 | 10505 | 1146 | 10.91 | 42 | 0.40 |
| 440 | 6082 | 879 | 14.45 | 34 | 0.56 |
| 700 | 8174 | 1610 | 19.70 | 93 | 1.14 |
| 950 | 16285 | 4019 | 24.68 | 783 | 4.80 |

**Table S3.** Relevant oligonucleotide sequences.

| Name | Sequence (5’-3’) |
| --- | --- |
| TSO | AAGCAGTGGTATCAACGCAGAGTGAATrGrGrG |
| IS | AAGCAGTGGTATCAACGCAGAGT |
| Adapter 1 | GATCGGAAGAGCACACGTCTGAACTCCAGTCAC |
| Adapter 2 | GCTCTTCCGATCT |
| P5-TSO primer | pAATGATACGGCGACCACCGAGATCTACACAAGCAGTGGTATCAACGCAGAGTAC |
| P7 primer | CAAGCAGAAGACGGCATACGAGATGTGACTGGAGTTCAGACGTGT |
| Splint oligo | GCCGTATCATTCAAGCAGAAGACG |

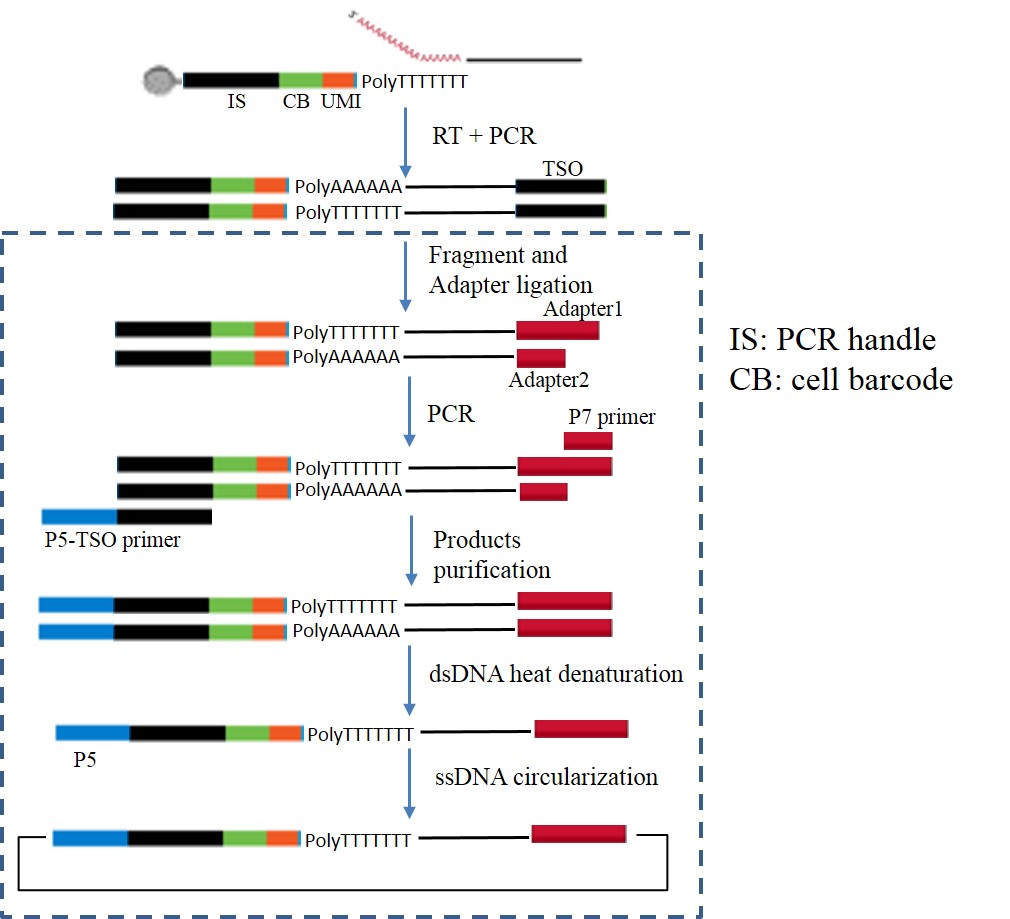

**Figure S1.** Library construction. The dashed box indicates the whole process of library construction.
